# Classification and characterisation of livestock production systems in northern Tanzania

**DOI:** 10.1101/2020.02.10.941617

**Authors:** W.A. de Glanville, A. Davis, K.J. Allan, J. Buza, J.R. Claxton, J.A. Crump, J.E.B. Halliday, P.C.D. Johnson, T.J. Kibona, B.T. Mmbaga, E.S. Swai, C. Uzzell, J. Yoder, J. Sharp, S. Cleaveland

## Abstract

Livestock keepers in sub-Saharan Africa face a growing range of pressures, including climate change, land loss, restrictive policies, and population increase. Widespread adaptation in response to such pressures can lead to the emergence of new, non-traditional typologies of livestock production.

We sought to characterise livestock production systems in northern Tanzania, a region undergoing rapid social, economic, and environmental change. Questionnaire and spatial data were collected from 404 livestock-keeping households in 21 villages in Arusha and Manyara Regions in 2016. Multiple factor analysis and hierarchical cluster analysis were used to classify households into livestock production systems based on household-level characteristics. Indicators of vulnerability, including household-level reports of hunger, illness, livestock loss, land loss and crop losses were compared between production systems.

Three distinct clusters emerged through this process. The ethnic, environmental and livestock management characteristics of households in each cluster broadly mapped onto traditional definitions of ‘pastoral’, ‘agro-pastoral’ and ‘smallholder’ livestock production in the region, suggesting that this quantitative classification system is complementary to more qualitative classification methods. Our findings also suggest that traditional systems of livestock production continue to persist in northern Tanzania. Nonetheless, we found indicators of substantial change within livestock production systems, most notably the adoption of crop agriculture in the majority of pastoral households. Smallholder households were less likely than either pastoral or agro-pastoral households to report hunger, illness, and livestock, land or crop losses.

Livelihoods that rely solely on livestock are relatively rare in northern Tanzania, which represents an important shift in production in the region, particularly among pastoralists. Policy initiatives to improve household and community well-being should recognise the continuing distinctiveness of traditional livestock production systems in the region.

## Introduction

Livestock play a key role in the livelihoods of many households in low-income countries. In Tanzania, 50% of all households keep livestock, with the sale of products derived from animals constituting an average of 15% of the annual income of rural livestock-keeping households[1]. Livestock in these settings also make important, but often under-recognised, contributions to livelihoods, for example as a basis for informal household insurance and financing, soil fertility, and labour saving, as well as to household nutrition through the production of animal source foods[2]. Indeed, livestock provide the social, cultural, and economic backbone to many rural economies in low-income settings, particularly those in marginal, semi-arid and arid environments. Here, the mobility of cattle, sheep, goats, and/or camels allows livestock keepers to utilise grazing and browsing on common land over a potentially wide geographic area[3], optimising production and reducing vulnerability to the effects of local rainfall deficits[4]. In these environments, livestock can also provide the security to pursue potentially riskier activities that rely on local rainfall, such as crop agriculture[4]. Supporting livestock production among the rural poor can provide an important route toward sustainable development, equitable livelihoods, and household health and welfare[5].

Livestock-based livelihoods are under growing pressure in many low-income countries from a range of sources[6]. These include the effects of climate change which, in East Africa, are expected to include increasing variability in precipitation[7–9]. Such effects are already becoming apparent in the region. In grassland areas of northern Tanzania, for example, the growing season during the ‘long rain’ period has declined from an average of 100 days in 1960 to 63 days in 2010[10]. Droughts in East Africa are also becoming more frequent and severe. In 2009, during one of the most severe droughts in living memory, up to 90% of livestock in some areas of northern Tanzania died[11]. Changing systems of land tenure, including the conversion of previously communal land to private ownership or wildlife conservation, further contribute to reduced availability of grazing land[12–16]. Livestock keepers in East Africa are therefore having to adapt to rapidly changing circumstances. Examples of adaptation include the adoption of non-traditional livestock species[17,18], new ways of rearing livestock[19], and the diversification of livelihood profiles in semi-arid areas away from livestock-focused production (i.e. pastoralism) toward mixed livestock and crop agriculture[20,21]. The extent of these changes and their implications for the characteristics and distribution of ‘traditional’ systems of livestock production in countries undergoing rapid social, economic, and environmental change warrants continued examination.

In northern Tanzania, three traditional typologies of livestock production (or livestock production systems) have existed for several centuries[22,23]. These systems of production can broadly be described as ‘pastoral’, ‘smallholder’, and ‘agro-pastoral’. While there has been substantial geographic and social overlap between systems, and their boundaries often hard to define[22], each has traditionally been linked to particular environmental conditions and ethnic groups. Pastoral systems have been found in the semi-arid, rangeland areas of northern Tanzania and historically dominated by Maasai ethnicities, with less populous groups such as the Barabaig also present. This production system has traditionally relied primarily (but not exclusively[22]) on livestock production, utilising long distance movements in response to variable rainfall patterns in an agriculturally marginal environment as a dominant risk-management strategy. Complex social organisation and systems of mutual support in response to the wide range of potential hazards that are present in these environments (including frequent droughts and livestock disease) have long been a feature of these communities[22]. Smallholder farming systems, by contrast, have traditionally been found on the high soil fertility slopes of Mount Kilimanjaro, Mount Meru and the Pare mountains. Here, members of ethnicities such as the Chagga, Meru, and Pare have reared typically small numbers of livestock that are integrated closely with intensive cash and subsistence crop production[23–25]. Agro-pastoral systems in northern Tanzania have also traditionally involved mixed crop and livestock agriculture but have typically been found in more marginal areas. While crop production has generally made the largest overall contribution to household livelihoods[26], large herd sizes with varying levels of mobility have allowed agro-pastoral farmers to also maximise the productivity of available grassland areas[4,27]. Agro-pastoral production in the region has historically been practiced by groups such as the Arusha and Iraqw, with the former having maintained particularly close social, cultural, and economic relationships with pastoral communities[23,28].

In light of livestock keeper adaptation to changing conditions in northern Tanzania, it is uncertain the extent to which these three broad typologies still characterise livestock production systems in the region. An evaluation of current characteristics of livestock production, and the classification of the production systems that exist in northern Tanzania, can contribute to the design of system-specific programmes that can support a range of livestock-based livelihoods[6,29]. It can also provide the basis for monitoring further change in these systems[6,29,30] and for identifying vulnerabilities to current and future hazards.

Myriad new livestock production typologies could emerge from demographic, technological, and environmental change. For example, a relatively small number of livestock keepers in Tanzania have adopted exclusively commercial production to meet growing demand for livestock products, particularly among urban populations. This has included beef ranching and the establishment of zero-grazing dairy units with European breeds of cattle[1]. The commercialisation and intensification of livestock production is strongly promoted by the Government of Tanzania[31]. Non-traditional, intensive production systems that have a greater focus on narrowly defined production objectives rather than subsistence or the socio-cultural utility of livestock are therefore likely to continue to emerge.

In addition, new technologies such as mobile telephones[32], new strategies and tools for household health management[33], and changes in land tenure and land availability[34] may lead to more subtle changes within traditionally defined production systems. While such adaptive change may increase overall diversity within a particular geographic area, it could also lead to further blurring of the boundaries between production systems. For example, the adoption of crop agriculture by Maasai pastoralists has been reported as a response to changing land tenure practices in northern Tanzania[20,21]. Widespread adoption and subsequent change within this traditional pastoral system could therefore conceivably lead to it becoming broadly indistinguishable (in terms of production) from neighbouring agro-pastoral systems.

Here, we use data generated from a cross-sectional survey of livestock-keeping households in northern Tanzania to classify and characterise livestock production systems at the household level in the region. Our main aim is to determine whether the three traditional typologies of livestock production (i.e., pastoral, agro-pastoral, smallholder) persist in northern Tanzania, and whether new systems of production can also be identified in the data. We explore variation in livestock production typologies in northern Tanzania across various dimensions, including ethnic and administrative boundaries. We use this analysis to consider how livestock production has changed in the region, how it may continue to change, and the implications of this change on the resilience of livestock keeping communities in northern Tanzania.

## Methods

### Study area

This work was conducted as part of the ‘Social, Environmental and Economic Drivers of Zoonotic disease’ (SEEDZ) project, a large cross-sectional study focused on human and animal zoonotic disease risk in six contiguous districts in Arusha Region (Arusha, Karatu, Longido, Meru, Monduli, and Ngorongoro Districts) and four contiguous districts in neighbouring Manyara Region (Babati Rural, Babati Urban, Mbulu, and Simanjiro Districts). Arusha and Manyara Regions are home to approximately 16% of all cattle and 26% of all sheep and goats in Tanzania[35,36]. The total human population is 3,119,441 in an area of 66,461 km^2^. The study area is made up of a mixture of semi-arid and sub-tropical agro-ecological zones[29].

### Village selection

Households were the unit of interest, with a multistage sampling design used to select them from within villages. Villages were selected using a generalised random tessalation stratified sampling (GRTS) approach, which provides a spatially balanced, probability-based sample[37]. The GRTS was performed using the *spsurvey* package[38] in the R statistical environment, version 3.1.1. (http://cran.r-project.org/). Village selection was made from a list of villages compiled from the 2012 National Census (Tanzanian National Bureau of Statistics, NBS). Villages in wards (an administrative unit comprising an average of 10 villages in the study area) that were classified as ‘urban’ rather than ‘rural’ or ‘mixed’ (i.e., urban and rural) by the 2012 census were excluded from the selection procedure. Villages inside the Ngorongoro Conservation Area (NCA), a wildlife area in which people and their livestock are permitted to live but in which crop agriculture is prohibited, were also excluded. With these exclusions, there were a total of 553 villages from which selection was made. To ensure sampling across a range of agro-ecological settings, villages in the study area were classified as those in which livestock-rearing (rather than crop agriculture) was considered to be the primary livelihood activity (‘pastoral’ villages) and those in which a mix of crop and livestock were considered as important (‘mixed’ villages). Village classification was performed in consultation with district-level government officials, typically the District Veterinary Officer or District Livestock Officer. Village selection was then stratified based on agro-ecological classifications, with 11 villages selected from those defined primarily as ‘pastoral’ and nine villages from those defined as ‘mixed.’ An additional village in a mixed setting was also selected non-randomly near our field headquarters on the outskirts of the city of Arusha for field trialling. No substantial changes were made to data collection tools after trialling, and we therefore include data collected from households in this village in this analysis.

Figure 1 shows the location of study villages in relation to the main landcover types in northern Tanzania.

**Figure 1.**
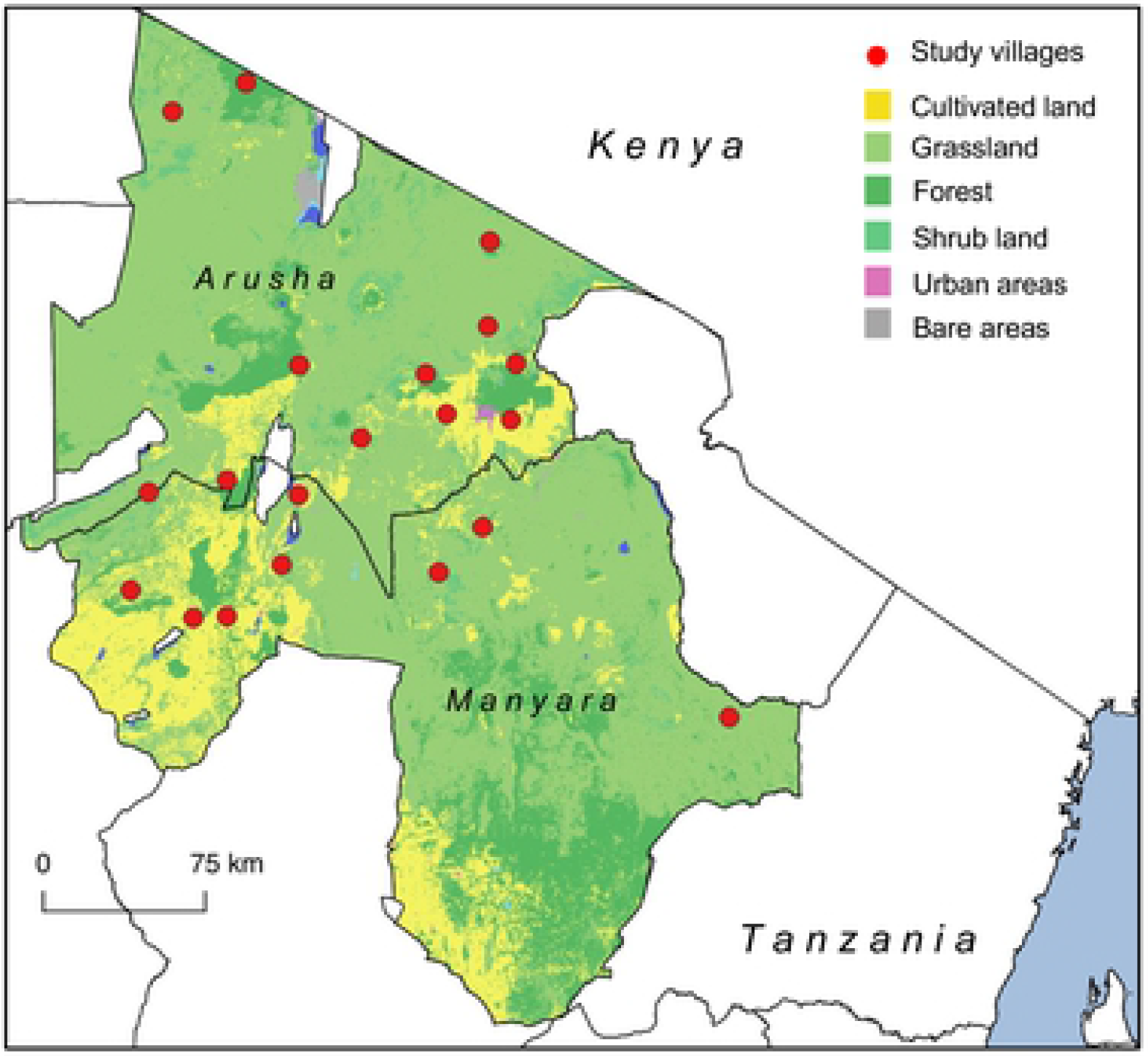
Map of study area in northern Tanzania showing location of study villages in relation to main land classifications in Arusha and Manyara Regions (Map created using QGIS version 2.14.3. Shape files from GADM; landcover raster data from Landsat (http://glcf.umd.edu/data/landsat/)).

### Household surveys

Study villages comprised between two and four sub-villages from which two or three were randomly selected for inclusion in the study. Within each sub-village, we adopted a central point sampling approach in which livestock keepers were invited to bring their animals to a pre-selected point within the sub-village, typically a livestock crush or dip tank. Data collection took place alongside sub-village level disease control activities, such as tick or worm control, conducted in collaboration with representatives from the Tanzanian Ministry of Livestock and Fisheries. Village authorities were notified of the proposed event at least three days in advance, with advertisement to livestock keepers in each sub-village made through the existing village administrative network of chairperson and village elders. During the sampling event, a list of all attending households was generated, and a maximum of ten households were selected from this list using a random number generator. During the central point event, we collected blood samples from animals owned by these households to test for infectious disease exposure, the results of which have been described elsewhere[39,40]. On a subsequent day, typically within one week, selected households were revisited. The household head received an in-depth questionnaire administered in either Kiswahili, Maa, or other local language by trained interviewers. The questionnaire covered a wide-range of topics, including household demographics, economics, livestock management practices and livestock health. The geographic co-ordinates of the household were captured using a handheld GPS (Garmin eTrex, Garmin Ltd, Olathe, Kansas, USA). Data collection took place between February and December 2016.

### Ethical approval

All participants provided written informed consent. The protocols, questionnaire tools and consent and assent procedures were approved by the ethics review committees of the Kilimanjaro Christian Medical Centre (KCMC/832) and National Institute of Medical Research (NIMR/2028) in Tanzania, and in the UK by the ethics review committee of the College of Medical, Veterinary and Life Sciences at the University of Glasgow (39a/15). Approval for study activities was also provided by the Tanzanian Commission for Science and Technology (COSTECH) and by the Tanzanian Ministry of Livestock and Fisheries, as well as by regional, district, ward and village-level authorities in the study area.

### Classification of livestock production systems

We used a data-driven approach to classify households into livestock production systems, which we define here as groups of households sharing the same or similar production characteristics[41]. Classification followed two stages. First, we performed dimension reduction using multiple factor analysis (MFA) on a selection of characteristics considered to represent variation between livestock-keeping households in the study area. Second, hierarchical cluster analysis (HCA) was performed on the output from the MFA (i.e. on a set of uncorrelated variables) with households grouped such that the within-group variability in household characteristics was minimized while between-group variability was maximized. The resulting clusters of households were interpreted to represent distinct and distinguishable livestock production system categories present in the study area at the time of the study.

### Dimension reduction by Multiple Factor Analysis (MFA)

Dimension reduction allows the variability among a set of potentially correlated variables to be represented in terms of a smaller, more parsimonious set of uncorrelated variables. Multiple factor analysis provides a dimension reduction approach for a set of variables describing categorical or continuous data that can be grouped in a meaningful way[42]. Eight groups of variables representing the characteristics of livestock keeping households in northern Tanzania were identified for use in the MFA procedure. The variable groupings (or domains) were: 1. Local household environment; 2. Household demographics; 3. Crop agriculture; 4. Numbers of cattle, sheep and goats owned; 5. Other livestock owned; 6. Livestock management practices; 7. Household food consumption practices; and 8. Indicators of household vulnerability. The variables comprising each of these domains are shown in Table 1. The MFA was performed in R using the *FactoMineR* package[43]. Data for household characteristics were derived from the household questionnaire (domains 2 to 8) or from data extracted at the household level within a geographic information system (QGIS, version 2.14.3) from publicly available environmental datasets (domain 1). Details on the source of environmental data and the questions asked at the household level are provided in the Supplementary Materials. Up to a maximum of 5% missing-ness was present in around 20% of variables. Imputation of missing values was performed using a regularized iterative MFA algorithm in the *missMDA* package[44] in R. Continuous variables (Domains 1 and 4) with obvious right skew were transformed using a natural logarithm. All continuous variables were scaled to have a mean of zero and a standard deviation of one before performing the MFA.

**Table 1.**
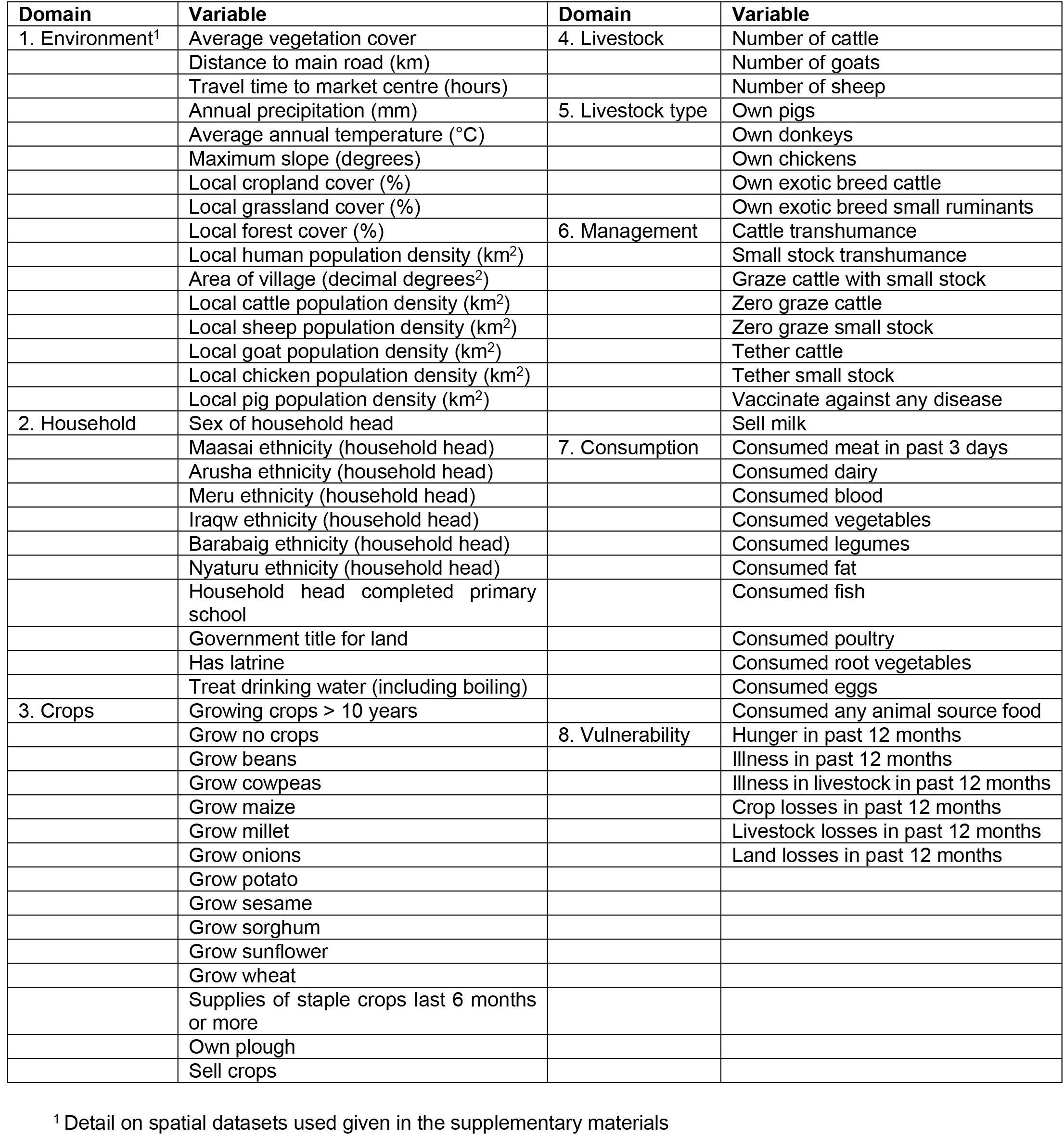
Domains and contributing variables for the multiple factor analysis to classify livestock-keeping households into production systems in northern Tanzania.

### Hierarchical cluster analysis (HCA)

Households were classified into clusters using HCA on the factors (i.e. the set of uncorrelated variables) derived from the MFA. To select which factors to include in the HCA, eigenvalues associated with each factor (describing how much variance is explained) were identified as ‘large’ or ‘small’ based on the presence of a natural break when consecutive eigenvalues were plotted on a scree plot[41]. All factors associated with ‘large’ eigenvalues were included in the clustering procedure. Ward’s minimum variance criteria were used to derive clusters, with no specification of the number of clusters made *a priori*. Hierarchical cluster analysis was performed using the *FactoMineR* package[43] in R. The average value of each household characteristic in each of the resulting clusters was compared to the global mean for that characteristic using the v-test[45]. A v-test value greater than 1.96 provides evidence (i.e. p-value <0.05) for a difference in the mean of the variable in the cluster when compared to the population mean.

## Results

Household survey data were collected from 404 households. The median (range) number of households interviewed per village was 19 (7, 30). Summary statistics for the household characteristics in each domain are given in Table 2 and 3.

**Table 2.** Mean values for continuous variables for households within clusters derived from hierarchical cluster analysis performed on livestock-keeping households in northern Tanzania (median values are given in square brackets).

| Domains | Variable | Mean [Median] |  |  |  |
| --- | --- | --- | --- | --- | --- |
|  |  | Overall<br>(n = 404) | Cluster 1<br>(n = 171) | Cluster 2<br>(n = 177) | Cluster 3<br>(n = 56) |
| Location | Average annual vegetation cover | 0.26 [0.27] | 0.23 [0.23] | 0.29 [0.29] | 0.27 [0.28] |
|  | Distance to main road (km) | 36.2 [32.6] | 47.8 [10.3] | 24.8 [9.4] | 7.3 [8.4] |
|  | Time to travel to market centre (hours) | 5.4 [3.6] | 6.4 [4.6] | 5.3 [3.4] | 2.6 [1.2] |
|  | Total annual precipitation (mm) | 830.4 [818] | 742.2 [741] | 865.6 [831] | 989.9 [912] |
|  | Average annual temperature (°C) | 19.3 [19.3] | 20.2 [20.0] | 18.6 [18.0] | 18.6 [18.6] |
|  | Maximum slope (degrees) | 3.7 [2.8] | 2.2 [1.5] | 4.5 [3.3] | 4.4 [4.0] |
|  | Local crop land cover (%) | 37.6 [25.7] | 9.5 [0.00] | 51.8 [49.7] | 78.3 [92.2] |
|  | Local grassland cover (%) | 46.1 [43.1] | 73.3 [89.5] | 31.1 [24.8] | 10.53 [1.2] |
|  | Local forest cover (%) | 1.1 [1.3] | 9.01 [0.14] | 14.29 [5.4] | 8.03 [0.58] |
|  | Local human population density (km<sup>2</sup>) | 1.41 [0.70] | 0.32 [0.17] | 1.28 [1.1] | 4.72 [1.1] |
|  | Area of village (decimal degrees<sup>2</sup>) | 1.8 [0.01] | 0.03 [0.03] | 0.01 [0.00] | 0.00 [0.00] |
|  | Local cattle density (km<sup>2</sup>) | 125.7 [0.90] | 2.4 [0.45] | 39.5 [9.7] | 784.3 [80.8] |
|  | Local sheep density (km<sup>2</sup>) | 95.4 [4.6] | 3.16 [2.7] | 10.31 [10.4] | 654.69 [17.1] |
|  | Local goat density (km<sup>2</sup>) | 61.6 [1.7] | 2.6 [1.27] | 17.5 [5.8] | 385.4 [41.7] |
|  | Local chicken density (km<sup>2</sup>) | 159.9 [9.4] | 4.9 [3.04] | 42.7 [13.3] | 1015 [155.0] |
|  | Local pig density (km<sup>2</sup>) | 2.0 [0.1] | 0.10 [0.02] | 1.1 [0.21] | 11.2 [0.09] |
| Livestock | Number cattle | 49.6 [10.0] | 104.5 [50.0] | 10.3 [8.0] | 6.5 [5.5] |
|  | Number goats | 52.6 [15.0] | 107.9 [50.0] | 13.6 [8.0] | 6.4 [4.5] |
|  | Number sheep | 50.0 [10.0] | 105.5 [42.0] | 8.4 [4.0] | 8.8 [3.5] |

### Multiple factor analysis

The percent contribution of each domain to explaining variation between households for the first two factors derived from the MFA is shown in Figure 2. The first factor (i.e. Dimension 1) explained 14.1% of the total variation, the second factor (Dimension 2) explained 6.3%, with all remaining factors each explaining less than 5%. The percent contribution to the inertia of the first factor was highest for Groups 1 (environment), 3 (crops), and 6 (livestock management) (Figure 2), reflecting the relative importance of these domains in explaining between-household variation.

**Figure 2.**
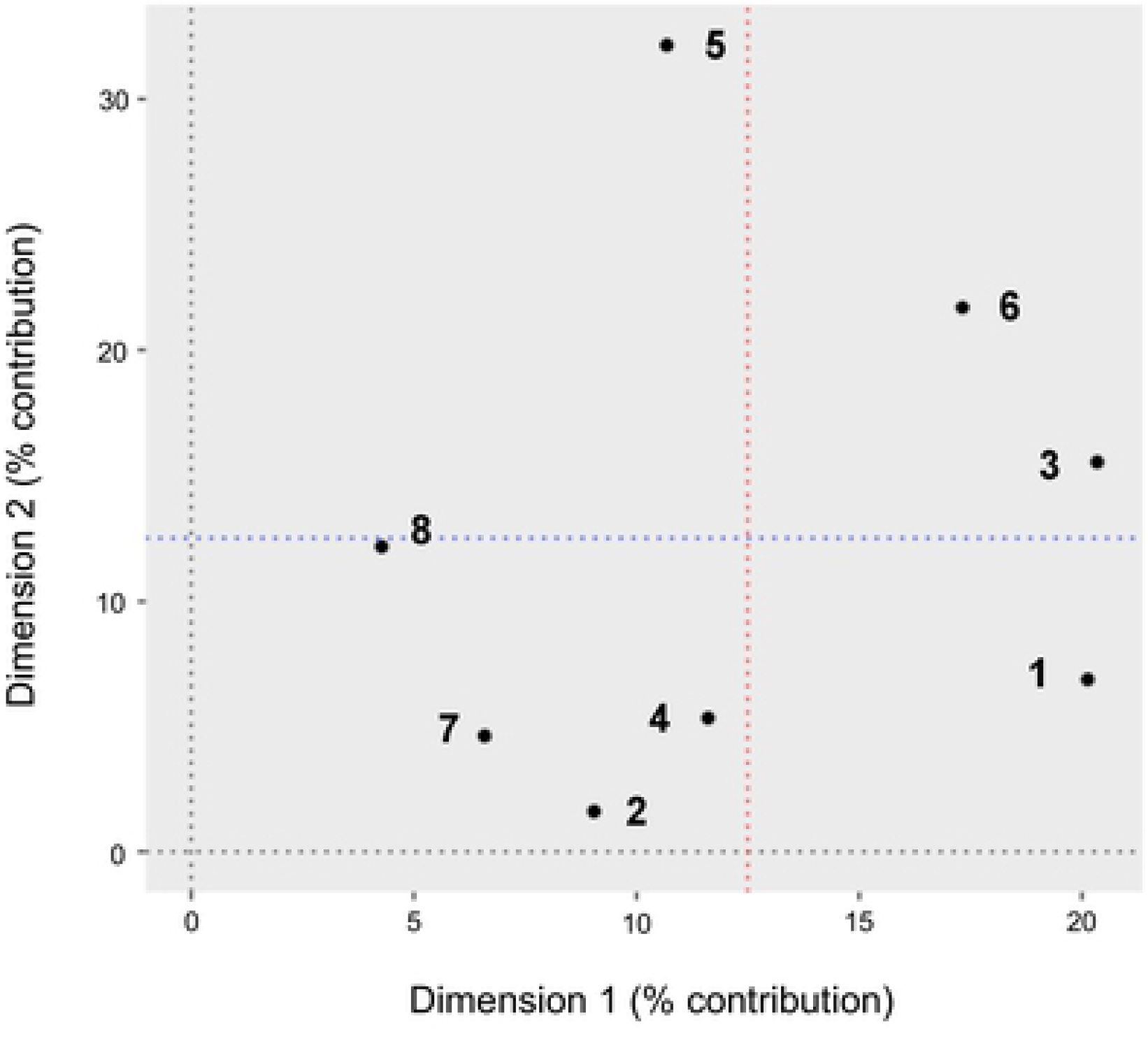
Percent contribution of each group to the first (dimension 1) and second (dimension 2) factors derived from MFA performed on characteristics of livestock-keeping households in northern Tanzania. Red (blue) dotted line represents the expected score if all domains contributed equally to the inertia on the first (second) factor (i.e. 100/8 = 12.5%).

Figure 3 shows the scores of those variables that made a contribution to the inertia of the first factor of greater than 1%. The average (median) contribution for included variables was 0.6% (0.2). The four categorical variables making the greatest overall contribution to the first factor were Maasai ethnicity of the household head (4.5%), not keeping donkeys (3.8%), engaging in cattle transhumance (3.4%), and engaging in small ruminant transhumance (3.2%). For the continuous characteristics, the top four variables were number of goats owned by a household (3.9%), number of cattle (3.0%), geographic area of village (2.5%), and human population density (2.5%). A full breakdown of all variable scores and their contributions to the first and second factors is given in the Supplementary Materials.

**Figure 3.**
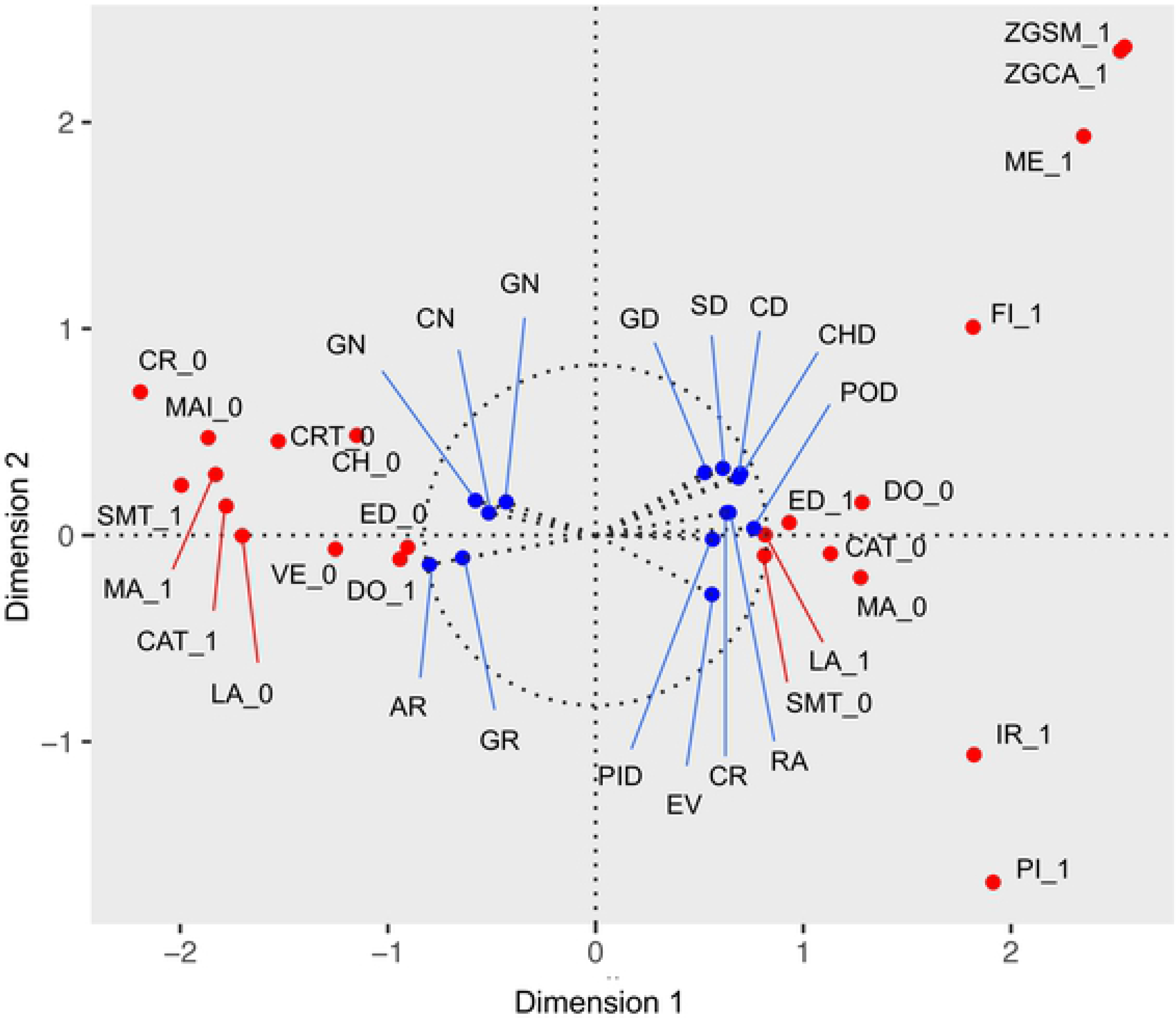
Variable scores in relation to the first and second factors derived from MFA performed on characteristics of livestock keeping households in northern Tanzania. Scores given to categorical (continuous) variables are shown in red (blue). ***Categorical*** (1 indicates presence of described characteristic; 0 indicates absence): **CAT** = Keep cattle; **CH** = Keep chickens; **CR** = Household grows crops; **CRT** = Grow crops for > 10 years; **DO** = Keep donkeys; **ED** = Household education to primary school or above; **FI** = Household consumed fish in past 3 days; **GCSM** = Graze cattle with small ruminants; **IR** = Iraqw ethnicity; **LA** = Latrine in household; **ME** = Meru ethnicity; **MA** = Maasai ethnicity; **MAI** = Grow maize; **PI** = Keep pigs; **SMT** = Small ruminant transhumance; **VE** = Household consumed vegetables in past 3 days; **ZGCA** = Zero graze cattle; **ZGSM** = Zero graze small ruminants. ***Continuous***: **AR** = Village area; **CD** = Cattle density; **CHD** = Chicken density; **CN** = Household cattle number; **CR** = Local cropland % cover; **EV** = Enhanced vegetation index; **GD** = Goat density; **GN** = Household goat number; **GR** = Local grassland % cover; **PID** = Pig density; **POD** = Human population density; **RA** = Annual precipitation; **SD** = Sheep density; **SN** = Household sheep number

Some clustering in scores of the categorical variables derived from the MFA is visually apparent in Figure 3. This includes the grouping of scores for variables such as Maasai-headed households, households that do not grow crops, or which have not been growing crops for more than 10 years, households engaging in cattle and small ruminant transhumance, households keeping donkeys but not chickens, and households without a latrine or in which the head does not have primary education clustering around negative values for Factor 1 (i.e. Dimension 1) and low negative and positive values on Factor 2 (Dimension 2). Scores for variables such as not engaging in transhumance, owning a latrine, some formal education of the household head, not owning a donkey, and grazing cattle with small ruminants tended to cluster around positive values for Factor 1 and low negative and positive values for Factor 2. There was a smaller cluster of scores for Iraqw-headed and pig-keeping households around positive values on Factor 1 and negative values on Factor 2, and a cluster of scores for Meru-headed and zero grazing households around positive values for Factor 1 and 2.

### Hierarchical cluster analysis

The HCA procedure resulted in three distinct clusters. The overall score on Factor 1 and 2 for study households and their membership of each cluster is shown in Figure 4. On the basis of the scree plot, the first five factors were included in the clustering procedure (see Supplementary Materials). The composition of each cluster in terms of continuous characteristics is described in Table 2 and in terms of categorical characteristics in Table 3. All continuous and categorical variables had a v-test score of greater than 1.96. The major differences in household characteristics between clusters can be summarised as:

**Table 3.**
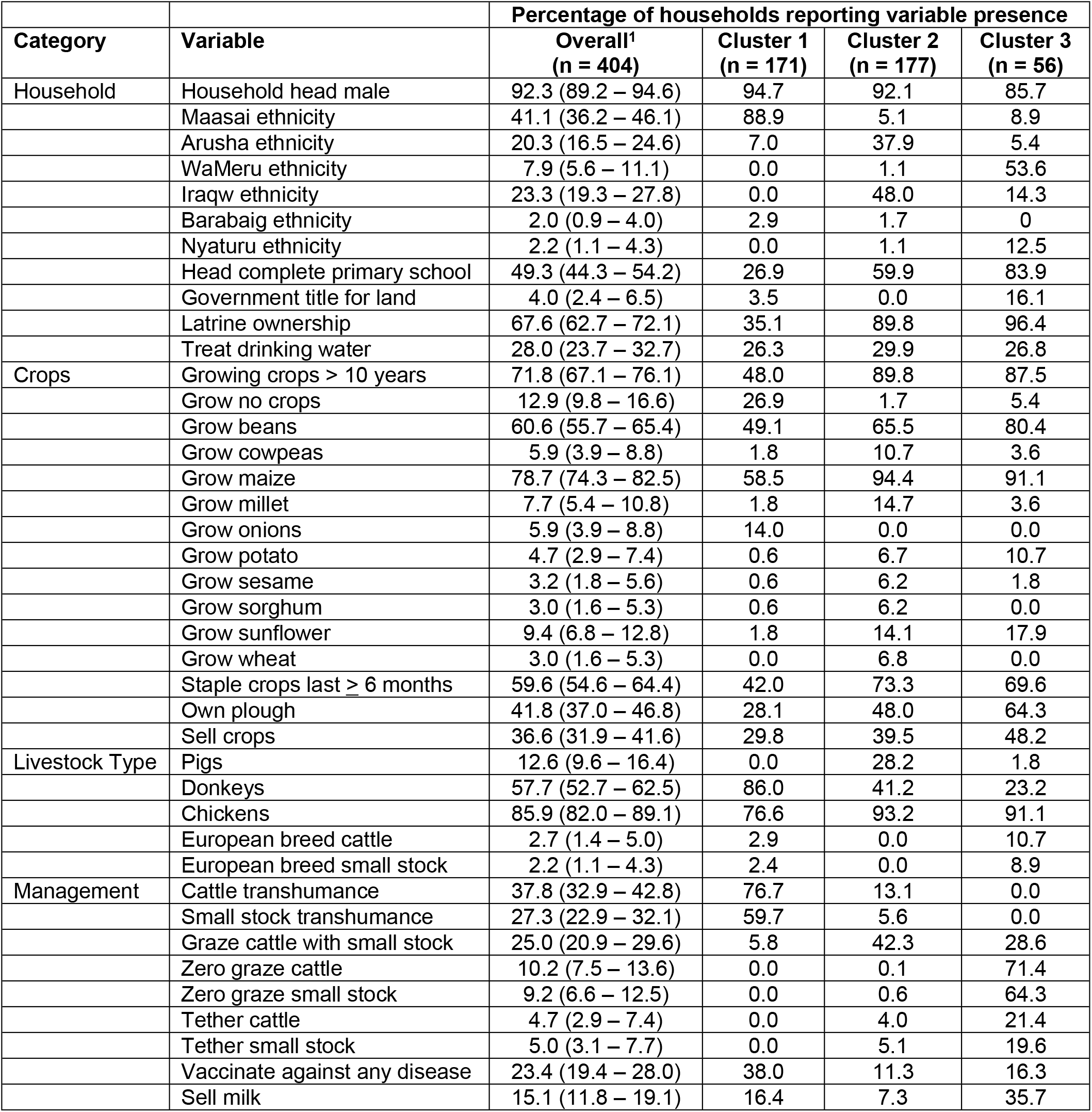
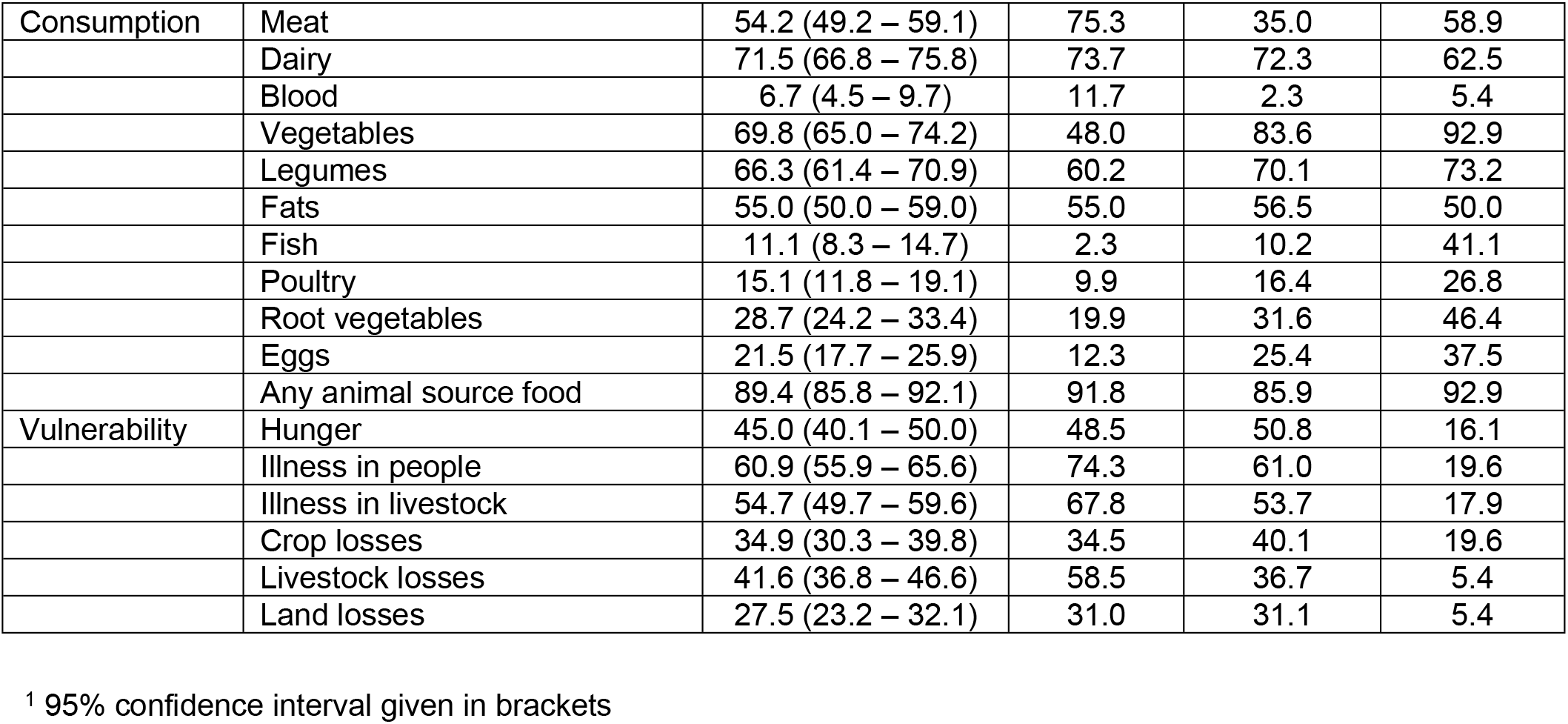
Percentages of households reporting variable presence in clusters derived from hierarchical cluster analysis performed on livestock-keeping households in northern Tanzania.

**Cluster 1**: Households in this cluster were characterised as being in areas with low average vegetation cover, having low levels of annual rainfall, low maximum slope (i.e., being relatively flat), low proportions of crop and forest land cover, and low population densities of both people and livestock. Cluster 1 households tended to be far from a main road and to have high average travel time to a market centre. Annual temperature, village area, and proportion of local grassland cover tended to be higher than for households in other clusters. Households in this cluster had the largest average herd sizes for cattle, sheep, and goats, and were typically headed by individuals with Maasai ethnicity, with 152 (91.6%) of 166 Maasai-headed households being found in this cluster. Other ethnicities found in this cluster included 12 (14.6%) of all 82 Arusha-headed households, 5 (63%) of the 8 Barabaig-headed households, and 1 (50%) of 2 Datoga-headed households. The majority of household heads in this cluster were without formal education beyond primary school and the proportion of households with a latrine was substantially lower than in the other two clusters. The majority of households reported growing crops in the past year, although this proportion was lower than in the other two clusters. Households growing onions were only found in this cluster. A relatively small proportion of households reported growing millet, sesame, or sunflower. A number of livestock management practices were commonly reported in this cluster, with households more commonly reporting transhumance for both cattle and small ruminants and using livestock vaccination in the past 12 months than households in the other two clusters. No households in this cluster reported zero grazing or using tethered grazing for cattle or small ruminants. No households in this cluster reported keeping pigs, but they commonly kept donkeys. Consuming meat in the past three days was commonly reported in this cluster. Household-level reports of illness in livestock and people, and reports of any livestock losses through mortality were also most common from households in this cluster. These reports were not adjusted for the number of people in a household or number of animals owned, the latter of which was highest in this cluster for all livestock species.

**Figure 4.**
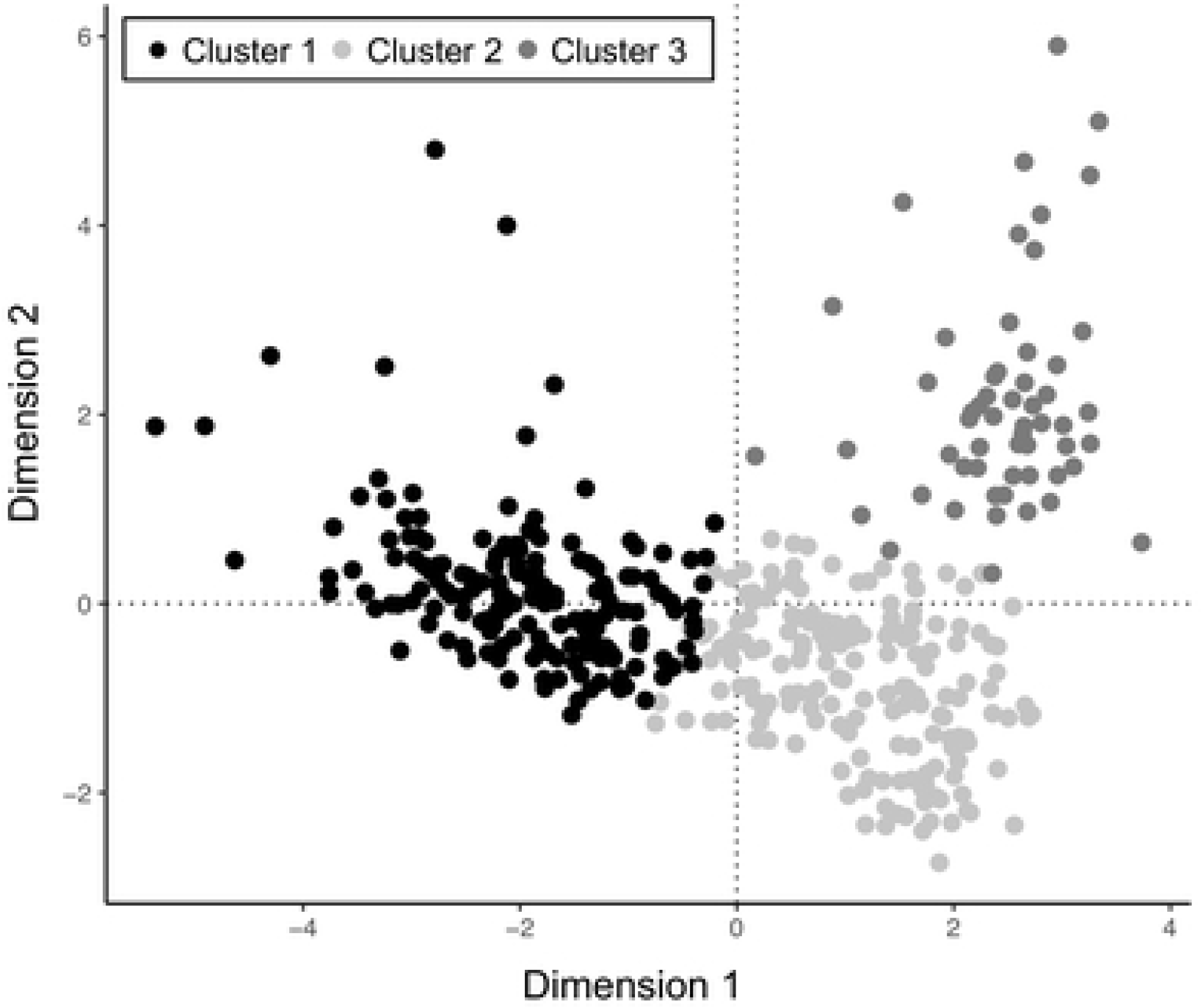
Position of households on the first and second factors (Dimension 1 and 2) derived from the MFA performed on characteristics of livestock keeping households in northern Tanzania. Households are shaded based on cluster membership.

**Cluster 2**: Households in this cluster typically had heads of Arusha and Iraqw ethnicity, including 67 (81.7%) out of 82 and 85 (90.4%) out of 94 of all households with heads with those ethnicities, respectively. Other ethnicities making up this cluster included 9 (5.4%) of the 166 Maasai-headed households, 3 (38%) out of 8 Barabaig-headed households, 1 out of 2 Datoga-headed households, 2 (6.3%) out of 32 Meru-headed households, 1 out of 2 Nyiramba-headed households and 2 (22.2%) out of 9 Nyaturu-headed households. All of the Burunge-(1), Luguru-(1), Rangi-(1), Sandawe-(3), and Sukuma-(1) headed households were in this cluster.

The mean, median and percentage values of most contributing variables in this cluster of households tended to fall between those for Clusters 1 and 3, with some exceptions. This cluster of households tended to be in areas with higher average vegetation cover and higher average proportion of local forest cover than those in Clusters 1 and 3. Households in this cluster were less likely than those in the other two clusters to have a government title for their land. Most households in this cluster reported growing crops in the past 12 months, with households growing cowpeas, millet, sesame, sorghum, and wheat most likely to be found in this cluster, as were households owning pigs and co-grazing cattle with small stock. Levels of livestock vaccination against any disease were lowest in this cluster. Households in this cluster were found in areas with the highest median pig population density. They were least likely to report consuming meat over the past 3 days. This was the largest cluster (Table 2).

**Cluster 3**: Households in this cluster tended to be closer to a main road and to have lower time to travel to a market centre than those in the other two clusters. They were in areas with relatively high annual rainfall, were most likely to be surrounded by cropland and least likely to be surrounded by grassland. Households in this cluster tended to be found in areas with the highest human, cattle, sheep, goat, and chicken population densities. They had the smallest cattle herd and goat flock sizes, but with average and median sheep flock sizes broadly equivalent with those in Cluster 2. Household heads in this cluster were most likely to be Meru ethnicity, including 30 (94%) out of all 32 Meru-headed households. Eight (8.5%) out of the 94 Iraqw-headed households, 5 (3.0%) out of the 166 Maasai-, 1 out of 2 Nyiramba-, and 7 (77.8%) out of 9 Nyaturu-headed households were also in this cluster. All of the Hehe-(1) and Chagga-(1) headed households were in this cluster. The proportion of households with heads with at least primary school education was highest in this cluster, as was the proportion of households with a latrine. The majority of households in the cluster reported growing crops, with the proportion of households growing beans, potatoes, sunflower and owning their own plough and reporting selling crops highest in this cluster. Relatively few households in this cluster reported owning donkeys. Ownership of exotic breed cattle and small ruminants was more commonly reported than in the other two clusters. No households in this cluster reported engaging in transhumance. Zero grazing cattle and small stock was common, as was tethering livestock for grazing. Households in this cluster most commonly reported consuming fish in the past three days. They were least likely to report hunger, illness in people or livestock, deaths in livestock, crop losses or land loss over the past 12 months. This was the smallest cluster (Table 2).

The proportion of households in each village assigned to each of these three clusters is shown in Figure 5. In seven villages, all households were members of Cluster 1; in three villages, all households were members of Cluster 2; and in one village, all households were members of Cluster 3. The remaining 10 study villages comprised a mixture of households from different clusters. Two villages had a mixture of households from all three clusters. When household cluster membership was compared to ‘pastoral’ village membership from the study design stage, 170 (82.9%) households in pastoral villages were in Cluster 1, 34 (16.6%) were in Cluster 2 and 1 (0.5%) was in Cluster 3. When compared to households in ‘mixed’ villages, 1 (0.5%) was in Cluster 1, 143 (71.9%) were in Cluster 2 and 55 (27.6%) were in Cluster 3.

**Figure 5.**
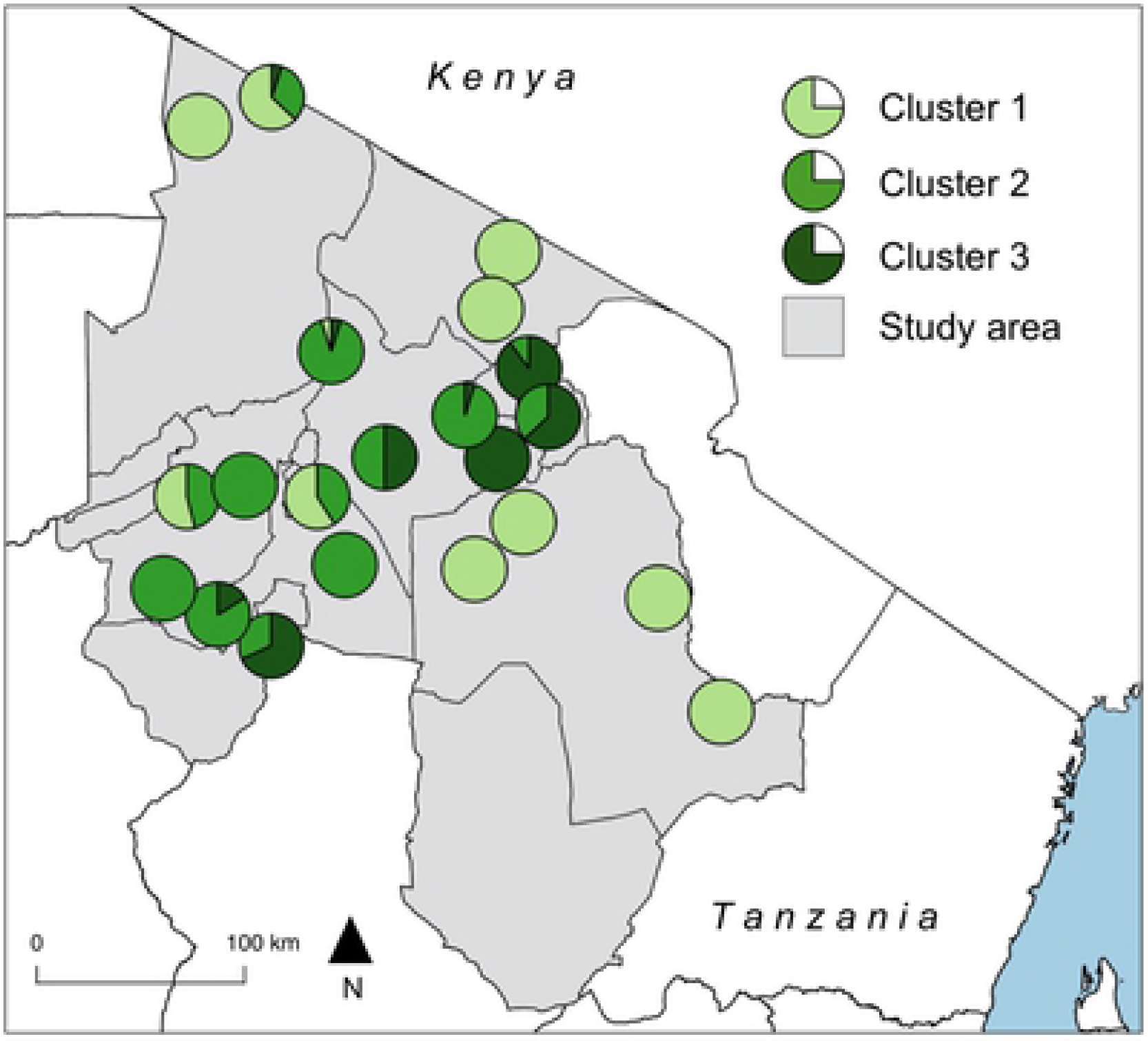
Proportion of households in study villages assigned to each livestock production cluster in northern Tanzania in 2016.

## Discussion

Our data analysis identified three clusters of households representing three distinct livestock production systems. The ethnic and production characteristics of these household clusters fit closely into the three traditional typologies of livestock production in northern Tanzania. These are pastoral (cluster 1), agro-pastoral (cluster 2) and smallholder (cluster 3) production systems. Our principal findings are therefore that the traditional livestock production systems that have existed in northern Tanzania for centuries continue to persist in the region, and that the analytical methods used herein complement more qualitative data categorization methods. While we find no evidence that new typologies of livestock production have emerged, there have been changes in production practices within existing systems. Our findings also reveal heterogeneity in a range of indicators of vulnerability between different production systems that point to inequalities in household health and welfare in northern Tanzania.

There has been a tendency, particularly reflected in livestock and land use policies, for pastoral communities to be viewed as static and resistant to change[46]. In reality, pastoral production systems are characterized by their ability to respond to highly changeable environments[47]. Here, we reveal widespread adoption of non-traditional forms of production within this system, most notably the fact that around three quarters of pastoral households reported performing crop agriculture over the preceding 12 months. Although crop agriculture has had an often under-appreciated role in East Africa pastoral livelihoods[22], the frequency with which crop agriculture was reported among pastoral households in this survey reflects a major shift in livelihoods, particularly for the Maasai[20,21]. An important driver for this change is likely to be the need to achieve greater food security as access to grazing lands declines, as well as to increase the security of land tenure, and provide access to additional sources of cash through the sale of crops such as maize and beans, the main crops reported to be grown by households in this group.

Mobility has also often been considered to be a defining characteristic of pastoral households[26]. It is therefore notable that over a quarter of cattle-keeping households in this cluster reported having not used transhumant grazing movements for cattle in the past 12 months, and more than one third of small ruminant keeping households of not using these movements for sheep or goats. It is well known that pastoral communities are undergoing rapid demographic, social, and economic shifts that are likely to influence practices around transhumance[48]. In particular, long-distance livestock movements that have traditionally been a response to variable grass and water availability have become increasingly difficult as a result of competing pressures on traditional grazing lands, including enclosure of previously communal land, conversion to crop lands, and for conservation [14,49–52]. It has also been argued that the rise of cultivation within pastoral systems may lead to reduced mobility and progression towards more sedentary systems in which livestock and crop agriculture are more closely integrated[53]. The impacts of restricted grazing and sedenterisation of pastoral communities have also been associated with declines in herd sizes in pastoral communities in other settings[54,55].

While our findings suggest the persistence of distinct pastoral and agro-pastoral livestock production systems in northern Tanzania, the ongoing ‘squeeze’ on rangeland access combined with growing populations is likely to lead to increasing overlaps in livestock production practices between these systems[47,56]. Strengthening extension services and the promotion of participatory initiatives that can support crop production in communities in which the cultural traditions of agriculture are relatively weak may be beneficial. It is notable, for example, that a very small proportion of pastoral households report growing indigenous crops such as sorghum or millet, which have relatively lower water requirements than introduced maize, and may represent less risky crop choices in dryland areas[57,58].

As livelihood transitions occur, it is also important to note that increasing reliance on crop agriculture in pastoral households can result in greater work for children, who may be expected to herd animals as well as work in fields, as well as for women, who perform most agricultural work in Maasai communities[52]. In addition, there is often limited integration of crop and livestock production in pastoral households, such as through the use of manure as fertiliser[47]. Unsustainable farming practices in combination with the common pastoral imperative to maximize herd sizes may also contribute to further rangeland declines if profits from agriculture are invested in additional livestock[4,47,59].

We find that livestock production systems in northern Tanzania are still strongly linked to ethnicity, but that these linkages are not absolute. For example, almost half of the Barabaig households in our sample were classified as being in the agro-pastoral livestock production system. This relatively small group of traditionally pastoral people is known to have been highly impacted by previous conversion of rangeland areas to commercial crop agriculture[60]. The long-term impacts of these changes have been infrequently assessed, but the results of our small sample of Barabaig-headed households may point to important changes in livelihood profiles away from pastoralism and towards agro-pastoralism. These changes may provide a model for a similar process underway in Maasai households. It is also striking that almost 10% of smallholder households were headed by people of Maasai ethnicity. Hence, modern Maasai households should be considered to include both pastoral and smallholder farmers, as well as those engaging in a wide range of non-livestock based livelihoods not considered here[61].

A notable finding in our study is the diversity of livestock production systems found within single villages. In many systems, be they social, ecological, or economic, increasing diversity tends to be correlated with increased resilience to a range of hazards[62]. It has been argued that the same is true of socio-ecological systems that are centred around livestock production[63]. For example, households that rear cattle, which are grazers, together with goats, which are browsers, enables the maximisation of livestock productivity under a range of environmental circumstances[63–65]. When systems of reciprocity within a single community are strong, multiple livestock-based livelihood strategies that allow different responses to hazards, such as drought or restrictive policies, might contribute to reducing the vulnerability of the whole community in a similar way. Systems of reciprocity have traditionally been an important feature of pastoral, agro-pastoral and smallholder production systems in northern Tanzania[66–68]. Such systems have been substantially eroded in recent times[14,69], and the extent to which they exist within administrative areas (such as villages) in which substantial diversity in livestock production typologies exist would be a valuable area for future research.

Our work reveals that the around half of all households reported hunger over the past 12 months in both pastoral and agro-pastoral production systems. While the proportion of pastoral households reporting consuming red meat, milk products, and blood over the past three days was equivalent to the other production systems, the proportion of households reporting consumption of vegetables, legumes, fish, eggs, and poultry was considerably lower. Dietary diversity has been strongly linked to food security[70] and to nutritional adequacy[71]. Household crop diversity was also low in all production systems, with maize and beans as the main crops grown. The production of a multiple crop types has been linked with lower levels of poverty[72,73].

The proportion of households with a government title for land was very low in all systems, and zero in the case of agro-pastoral households. One third of households in this group also report land losses in the past 12 months. Land insecurity is strongly linked to poverty vulnerability and can be expected to become an increasing issue with population growth in the region[49]. Efforts in pastoral communities have been made by local non-governmental organisations to facilitate the securing of land titles and land rights, but to mixed effect[34]. Household-level latrine ownership and the education of the household head were considerably less common in pastoral households than in the other production systems. These indicators of household socioeconomic status have been strongly linked to human infectious disease risk[74] and childhood nutrition[75–77] in other settings. In many pastoral communities, sedentarisation has been associated with negative nutritional and health consequences, despite often improved access to services[78].

The smallholder production system had the lowest proportion of households reporting hunger, illness in people or livestock, livestock or crop losses or land losses. This group also tended to report the widest diversity of food consumption and had the highest level of household head education and latrine ownership. While we did not collect detailed data on household inputs as part of this study, smallholder systems in northern Tanzania have historically represented very high levels of agricultural intensification[23]. The apparent resilience (or lower levels of vulnerability) of households within this system support links between agricultural intensification and prosperity[56], although this group was found in peri-urban areas and are therefore also likely to benefit from greater access to extension and other services, as well as non-agricultural sources of income that were not recorded here. Smallholder systems are often a focus for development interventions in the livestock sector in sub-Saharan Africa[80]. However, based on the indicators of vulnerability explored in this study, and given scarce resources, livestock-keeping households in agro-pastoral and pastoral settings in northern Tanzania appear to be in greater need of support in poverty alleviation.

A number of limitations should be considered when interpreting our findings. Households were selected from a limited number of villages, and villages in urban areas were excluded from the sampling procedure. Livestock production occurs in urban areas of Tanzania and tends to be characterised by small scale, intensive zero-grazing production of cattle and small ruminants that could be expected to fall into the smallholder classification. The proportion of households that were categorised as smallholder was smaller than those in the other systems. With a larger sample, greater diversity within the smallholder system may have emerged, potentially including the classification of distinct typologies. In particular, while zero grazing practices were common in the smallholder system, relatively few households reported ownership of European breed dairy cattle or the sale of milk. Hence, greater sampling in smallholder settings, including in villages classified as ‘urban’ may have revealed a distinct typology involving high yielding European breed cattle kept exclusively for commercial purposes. In addition, we collected only limited information on household livelihood activities outside of crop and livestock production, or the relative contribution of each to household revenues. While we included a wide range of household level characteristics, and the resulting clusters reflect expected and sensible groupings of livestock-keeping households with these characteristics in this region of Tanzania, dimension reduction and hierarchical clustering approaches are sensitive to input data. We therefore cannot rule out that the inclusion of a wider range of household level variables than were available to us may have resulted in a different number of clusters, or clusters with different general characteristics. A further limitation is that all livestock keeping households included in this study were those who attended the central point sampling event. There is therefore the potential for selection bias if characteristics of households made them more or less likely to attend with their animals. This may be particularly important for those households in the smallholder sector, where zero-grazing (i.e. continual housing) of animals was most commonly reported. Finally, while not necessarily a limitation, we would caution that the production systems we describe here represent those in northern Tanzania, and “smallholders”, “agro-pastoralists” and “pastoralists” may have different characteristics in other parts of the country and internationally. Future studies that use a similar approach to that described here to classify livestock production systems in other geographic areas would provide further understanding of the diversity of livestock production that exists in Tanzania.

Previously reported classification systems have commonly used knowledge-based systems, such as expert opinion, in order to classify large geographic areas according to the dominant livestock production system [4,30,81–84]. The resulting classification systems have made important contributions to priority-setting, but their regional, continental or global focus has meant that they typically have limited resolution at smaller spatial scales. Here we show that data-driven approaches performed on the types of data variables that are commonly collected in questionnaire-based surveys, can provide a valuable tool with which to characterise and classify livestock keeping households. We show that such an approach can allow the diversity of livestock production that can exist within small areas, including within a single village, to be described.

## Acknowledgements

We would like to thank the livestock-keepers who participated in this study, as well as village, ward, district and regional authorities for the facilitation of this work. We are very grateful to the SEEDZ field team including Kunda Mnzava, Tauta ole Maapi, Rigobert Tarimo, Fadhili Mshana, Zanuni Kweka, Euphrasia Mariki, Mamus Toima, Matayo Melubo, Sambeke Melubo, and Hassan Hussein for their contribution to data collection.

## Supporting Information

**S1 File.** Supplementary materials

## References

1. Covarrubias K, Nsiima L, Zezza A. Livestock and livelihoods in rural Tanzania: a descriptive analysis of the 2009 National Panel Survey. Joint Paper of the World Bank, FAO, AU-IBAR, ILRI and the Tanzanian Ministry of Livestock and Fisheries Development; 2012.

2. Herrero M, Grace D, Njuki J, Johnson N, Enahoro D, Silvestri S, et al. The roles of livestock in developing countries. Animal. 2013;7 Suppl 1: 3–18. doi:10.1017/S1751731112001954

3. Ogola J, Fèvre EM, Gitau GK, Christley R, Muchemi G, de Glanville WA. The topology of between-herd cattle contacts in a mixed farming production system in western Kenya. Prev Vet Med. 2018;158: 43–50. doi:10.1016/j.prevetmed.2018.06.010

4. Jahnke HE. Livestock Production Systems and Livestock Development in Tropical Africa. Kieler Wissenschaftsverlag Vauk. 273 pp. 1982.

5. FAO. World Livestock: Transforming the livestock sector through the Sustainable Development Goals. Food and Agriculture Organization of the United Nations. Rome. 12 pp. 2018.

6. Thornton PK. Livestock production: recent trends, future prospects. Philos Trans R Soc Lond, B, Biol Sci. 2010;365: 2853–2867. doi:10.1098/rstb.2010.0134

7. Shongwe ME, van Oldenborgh GJ, van den Hurk B, van Aalst M. Projected Changes in Mean and Extreme Precipitation in Africa under Global Warming. Part II: East Africa. Journal of Climate. 2011;24: 3718–3733. doi:10.1175/2010JCLI2883.1

8. Ongoma V, Chen H, Gao C. Projected changes in mean rainfall and temperature over East Africa based on CMIP5 models. International Journal of Climatology. 2018;38: 1375–1392. doi:10.1002/joc.5252

9. Tierney JE, Ummenhofer CC, de Menocal PB. Past and future rainfall in the Horn of Africa. Sci Adv. 2015;1: e1500682. doi:10.1126/sciadv.1500682

10. Kihupi N, Tarimo A, Masika R, Boman B, Dick W. Trend of growing season characteristics of semi-arid Arusha District in Tanzania. Journal of Agricultural Science. 2015;7: 45–55.

11. Goldman MJ, Riosmena F. Adaptive Capacity in Tanzanian Maasailand: Changing strategies to cope with drought in fragmented landscapes. Global Environmental Change. 2013;23: 588–597.

12. Nyariki DM, W Mwang’ombe A, Thompson DM. Land-Use Change and Livestock Production Challenges in an Integrated System: The Masai-Mara Ecosystem, Kenya. Journal of Human Ecology. 2017;26: 163–173. doi:10.1080/09709274.2009.11906178

13. Archambault CS. Re-creating the commons and re-configuring Maasai women’s roles on the rangelands in the face of fragmentation. International Journal of the Commons. 2016;10: 728–746.

14. Lesorogol CK. Land Privatization and Pastoralist Well-being in Kenya. Development & Change. 2008;39: 309–331. doi:10.1111/j.1467-7660.2007.00481.x

15. Shiferaw B, Tesfaye K, Kassie M, Abate T, Prasanna BM, Menkir A. Managing vulnerability to drought and enhancing livelihood resilience in sub-Saharan Africa: Technological, institutional and policy options. High Level Meeting on National Drought Policy. 2014;3: 67–79.

16. Okello MM. Land Use Changes and Human–Wildlife Conflicts in the Amboseli Area, Kenya. Human Dimensions of Wildlife. 2005;10: 19–28. doi:10.1080/10871200590904851

17. Kagunyu AW, Wanjohi J. Camel rearing replacing cattle production among the Borana community in Isiolo County of Northern Kenya, as climate variability bites. Pastoralism. 2014;4: 13.

18. Sperling L. The Adoption of camels by Samburu cattle herders. Nomad People. White Horse Press; 1987: 1–17.

19. Pretty J, Toulmin C, Williams S. Sustainable intensification in African agriculture. International Journal of Agricultural Sustainability. 2011;9: 5–24. doi:10.3763/ijas.2010.0583

20. McCabe JT, Leslie PW, Deluca L. Adopting Cultivation to Remain Pastoralists: The Diversification of Maasai Livelihoods in Northern Tanzania. Human Ecology. 2010;38: 321–334. doi:10.1007/s10745-010-9312-8

21. McCabe JT. Sustainability and Livelihood Diversification among Maasai of Northern Tanzania. Human Organization. 2003;62: 100–111.

22. Spear T, Waller R. Being Maasai: ethnicity and identity in East Africa. Ohio University Press; 1993.

23. Spear T. Mountain farmers: moral economics of land and agricultural development in Arusha and Meru. University of California Press, Berkeley; 1997. pp. 1–262.

24. Fernandes ECM, Oktingati J, Maghembe J. The Chagga homegardens: a multistoried agroforestry cropping system on Mt. Kilimanjaro (Northern Tanzania). Agroforestry Systems. 1984;2: 73–86.

25. Maro P. Population and land resources in northern Tanzania: the dynamics of change 1920 - 1970. PhD Thesis, Dept. of Geography, University of Minnesota. 1975.

26. Morton J, Meadows N. Pastoralism and Sustainable Livelihoods: An Emerging Agenda. Policy Series 11. National Resources Institute, University of Greenwich, 56 pp.

27. Otte M, Chilonda P. Cattle and small ruminant production systems in sub-Saharan Africa: a systematic review. Food and Agriculture Organization of the United Nations, Rome; 2002.

28. Ole Kuney RO. Pluralism and ethnic conflict in Tanzania’s arid lands: the case of the Maasai and the WaArusha. Nomad People. 1994; 95–107.

29. Robinson TP, Thornton PK, Franceschini G, Kruska RL, Chiozza F, Notenbaert A, et al. Global livestock production systems. Rome, Food and Agriculture Organization of the United Nations (FAO) and International Livestock Research Institute (ILRI), 152 pp. 2011.

30. Steinfeld H, Wassenaar T, Jutzi S. Livestock production systems in developing countries: status, drivers, trends. Rev Sci Tech. 2006;25: 505–516.

31. Ministry of Livestock and Fisheries Development. Livestock modernization initiative. United Republic of Tanzania, Ministry of Livestock and Fisheries Development. 2015.

32. Debsu DN, Little PD, Tiki W, Guagliardo SAJ, Kitron U. Mobile phones for mobile people: the role of information and communication technology (ICT) among livestock traders and Borana pastoralists of southern Ethiopia. Nomad People. 2016;20: 35–61.

33. Schelling E, Wyss K, Diguimbaye C, Béchir M, Taleb MO, Bonfoh B, et al. Towards Integrated and Adapted Health Services for Nomadic Pastoralists and their Animals: A North–South Partnership. In: Hadorn GH, Hoffmann-Riem H, Biber-Klemm S, Grossenbacher-Mansuy W, Joye D, Pohl C, et al., editors. Handbook of Transdisciplinary Research. Dordrecht: Springer Netherlands; 2008. pp. 277–291.

34. Goldman MJ, Davis A, Little J. Controlling land they call their own: access and women’s empowerment in Northern Tanzania. The Journal of Peasant Studies. 2015;43: 777–797. doi:10.1080/03066150.2015.1130701

35. National Bureau of Statistics. United Republic of Tanzania. National Sample Census of Agriculture 2002/2003: Arusha Region. 2007.

36. National Bureau of Statistics. United Republic of Tanzania. National Sample Census of Agriculture 2002/2003: Manyara Region. 2007.

37. Stevens DL, Olsen AR. Spatially Balanced Sampling of Natural Resources. J Am Stat Assoc. 2004;99: 262–278.

38. Kincaid TM, Olsen AR. spsurvey: Spatial Survey Design and Analysis. R package version 2.3. US Environmental Protection Agency. 2012.

39. Herzog CM, de Glanville WA, Willett BJ, Kibona TJ, Cattadori IM, Kapur V, et al. Pastoral production is associated with increased peste des petits ruminants seroprevalence in northern Tanzania across sheep, goats and cattle. Epidemiol Infect. 2019;147: e242. doi:10.1017/S0950268819001262

40. Semango G, Hamilton CM, Kreppel K, Katzer F, Kibona T, Lankester F, et al. The Sero-epidemiology of Neospora caninum in Cattle in Northern Tanzania. Front Vet Sci. 2019;6: 1473. doi:10.3389/fvets.2019.00327

41. Costard S, Porphyre V, Messad S, Rakotondrahanta S, Vidon H, Roger F, et al. Multivariate analysis of management and biosecurity practices in smallholder pig farms in Madagascar. Prev Vet Med. 2009;92: 199–209. doi:10.1016/j.prevetmed.2009.08.010

42. Abdi H, Williams LJ, Valentin D. Multiple factor analysis: principal component analysis for multitable and multiblock data sets. Comp Stat. 2013;5: 149–179.

43. Le S, Josse J, Husson F. FactoMineR: An R Package for Multivariate Analysis. J Stat Softw. 2008;25: 1–18.

44. Husson F, Josse J, Le S, J M. FactoMineR: Multivariate Exploratory Data Analysis and Data Mining with R. R Package Version 1.25. http://CRAN.R-project.org/package=FactoMineR. 2013.

45. Morineau A. Note sur la caracterisation statistique d’une classe et les valeurs tests. Bull Techn Centre Statist Inform Appl. 1984; 9–12.

46. De Carvalho EC. “Traditional” and “Modern” Patterns of Cattle Raising in Southwestern Angola: A Critical Evaluation of Change from Pastoralism to Ranching. The Journal of Developing Areas. 1974;8: 199–226.

47. Conroy AB. Maasai oxen, agriculture and land use change in Monduli District, Tanzania. PhD Thesis. University of New Hampshire. 2001.

48. Turner MD, Schlecht E. Livestock mobility in sub-Saharan Africa: A critical review. Pastoralism. 2019;9: 13.

49. Little PD, McPeak J, Barrett CB, Kristjanson P. Challenging Orthodoxies: Understanding Poverty in Pastoral Areas of East Africa. Development & Change. 2008;39: 587–611. doi:10.1111/j.1467-7660.2008.00497.x

50. Börjeson L, Hodgson DL, Yanda PZ. Northeast Tanzania’s Disappearing Rangelands: Historical Perspectives on Recent Land Use Change. The International Journal of African Historical Studies. 2008;41: 523–556.

51. Shiferaw B, Tesfaye K, Kassie M, Abate T, Prasanna BM, Menkir A. Managing vulnerability to drought and enhancing livelihood resilience in sub-Saharan Africa: Technological, institutional and policy options. Weather and Climate Extremes. 2014;3: 67–79.

52. Archambault C. Re-creating the commons and re-configuring Maasai women’s roles on the rangelands in the face of fragmentation. International Journal of the Commons. 2016;10: 728–746.

53. Bourn D, Wint W. Livestock, land-use and agricultural intensification in sub-Saharan Africa. London: Overseas Development Institute. 1994.

54. Tilahun M, Angassa A, Abebe A, Mengistu A. Perception and attitude of pastoralists on the use and conservation of rangeland resources in Afar Region, Ethiopia. Ecological Processes. 2016;5: 18.

55. Kimiti KS, Western D, Mbau JS, Wasonga OV. Impacts of long-term land-use changes on herd size and mobility among pastoral households in Amboseli ecosystem, Kenya. Ecological Processes. 2018;7: 4.

56. Boserup E. The Conditions of Agricultural Growth. Earthscan UK, Abingdon; 1965.

57. Omoyo NN, Wakhungu J, Oteng’i S. Effects of climate variability on maize yield in the arid and semi arid lands of lower eastern Kenya. Agriculture & Food Security. 2015;4: 8.

58. Karanja DR, Githunguri CM, MRagwa L, Mulwa D, Mwiti S. Variety characteristics and production guidelines of traditional food crops. Kenya Agricultural Research Institute.

59. Wynants M, Kelly C, Mtei K, Munishi L, Patrick A, Rabinovich A, et al. Drivers of increased soil erosion in East Africa’s agro-pastoral systems: changing interactions between the social, economic and natural domains. Regional Environmental Change. 2019;19: 1909–1921.

60. Lane CR. Pastures lost: alienation of Barabaig land in the context of land policy and legislation in Tanzania. Nomad People. 1994;: 81–94.

61. Hauck S, Rubenstein DI. Pastoralist societies in flux: A conceptual framework analysis of herding and land use among the Mukugodo Maasai of Kenya. Pastoralism. 2017;7: 18.

62. Elmqvist T, Folke C, Nyström M, Peterson G, Bengtsson J, Walker B, et al. Response diversity, ecosystem change, and resilience. Frontiers in Ecology and the Environment. 2003;1: 488–494. doi:10.1890/1540-9295(2003)001[0488:RDECAR]2.0.CO;2

63. Leslie P, McCabe JT. Response Diversity and Resilience in Social-Ecological Systems. Curr Anthropol. 2013;54: 114–143.

64. Mace R, Houston A. Pastoralist strategies for survival in unpredictable environments: A model of herd composition that maximises household viability. Agricultural Systems. 1989;31: 185–204.

65. Little PD, Smith K, Cellarius BA, Coppock DL, Barrett C. Avoiding Disaster: Diversification and Risk Management among East African Herders. Development & Change. 2001;32: 401–433. doi:10.1111/1467-7660.00211

66. Aktipis A, de Aguiar R, Flaherty A, Iyer P, Sonkoi D, Cronk L. Cooperation in an Uncertain World: For the Maasai of East Africa, Need-Based Transfers Outperform Account-Keeping in Volatile Environments. Human Ecology. 2016;44: 353–364.

67. Howard M. Socio-economic causes and cultural explanations of childhood malnutrition among the Chagga of Tanzania. Soc Sci Med. 1994;38: 239–251.

68. Rekdal OB. Money, Milk and Sorghum Beer: Change and Continuity among the Iraqw of Tanzania. Africa: Journal of the International African Institute. 1996;66: 367–385.

69. Howard M. Socio-economic causes and cultural explanations of childhood malnutrition among the Chagga of Tanzania. Soc Sci Med. 1994;38: 239–251.

70. Schwei RJ, Tesfay H, Asfaw F, Jogo W, Busse H. Household dietary diversity, vitamin A consumption and food security in rural Tigray, Ethiopia. Public Health Nutr. 2017;20: 1540–1547. doi:10.1017/S1368980017000350

71. Arimond M, Ruel MT. Dietary Diversity Is Associated with Child Nutritional Status: Evidence from 11 Demographic and Health Surveys. J Nutr. 2004;134: 2579–2585.

72. Birthal PS, Roy D, Negi DS. Assessing the Impact of Crop Diversification on Farm Poverty in India. World Development. 2015;72: 70–92.

73. Ecker O. Agricultural transformation and food and nutrition security in Ghana: Does farm production diversity (still) matter for household dietary diversity? Food Policy. 2018;79: 271–282.

74. de Glanville WA, Thomas LF, Cook EAJ, Bronsvoort BM de C, Wamae NC, Kariuki S, et al. Household socio-economic position and individual infectious disease risk in rural Kenya. Sci Rep. 2019;9: 2972. doi:10.1038/s41598-019-39375-z

75. Makoka D, Masibo PK. Is there a threshold level of maternal education sufficient to reduce child undernutrition? Evidence from Malawi, Tanzania and Zimbabwe. BMC Pediatr. 2015;15: 96. doi:10.1186/s12887-015-0406-8

76. Wachs TD, Creed-Kanashiro H, Cueto S, Jacoby E. Maternal education and intelligence predict offspring diet and nutritional status. J Nutr. 2005;135: 2179–2186. doi:10.1093/jn/135.9.2179

77. Boyle MH, Racine Y, Georgiades K, Snelling D, Hong S, Omariba W, et al. The influence of economic development level, household wealth and maternal education on child health in the developing world. Soc Sci Med. 2006;63: 2242–2254. doi:10.1016/j.socscimed.2006.04.034

78. Roth EA, Fratkin E. The Social, Health and Economic Consequences of Pastoral Sedentarization in Marsabit District, Northern Kenya. In: Fratkin E, Roth EA, editors. As Pastoralists Settle: Social, Health and Economic Consequences of Pastoral Sedentarization in Marsabit District, Kenya. Springer, Boston, MA; 2005. pp. 1–28.

79. Vohland K, Barry B. A review of in situ rainwater harvesting (RWH) practices modifying landscape functions in African drylands. Agriculture, Ecosystems and Environment. 2009;131: 119–127.

80. Bill and Melinda Gates Foundation. Agricultural development strategy overview. Available at: https://www.gatesfoundation.org/what-we-do/global-growth-and-opportunity/agricultural-development.

81. Thornton PK, Jones PJ, Owiyo TM, Kruska RL, Herrero M, Kristjanson P, et al. Mapping climate vulnerability and poverty in Africa. 200 pp. Nairobi, International Livestock Research Institute (ILRI). 2006 May.

82. Otte J, Chilonda P. Classification of Cattle and Small Ruminant Production Systems in Sub-Saharan Africa. Outlook on Agriculture. 2003;32.

83. Cecchi G, Wint W, Shaw A, Marletta A, Mattioli R, Robinson TP, et al. Geographic distribution and environmental characterization of livestock production systems in Eastern Africa. Agriculture, Ecosystems and Environment. 2010;135: 98–110.

84. Sere C, Steinfeld H. World livestock production systems. FAO Animal Production and Health Paper 127. Rome, Food and Agriculture Organization of the United Nations (FAO). 1996 Jun.

